# Identification of functionally connected multi-omic biomarkers for Alzheimer’s Disease using modularity-constrained Lasso

**DOI:** 10.1101/789693

**Authors:** Linhui Xie, Pradeep Varathan, Kwangsik Nho, Andrew J. Saykin, Paul Salama, Jingwen Yan

**Affiliations:** Department of Electrical and Computer Engineering, Indiana University Purdue University Indianapolis, Indianapolis, Indiana, USA; Department of Radiology and Imaging Sciences, School of Medicine, Indiana University School of Medicine, Indianapolis, Indiana, USA; Department of BioHealth Informatics, Indiana University Purdue University Indianapolis, Indianapolis, Indiana, USA

## Abstract

In the past decade, a large number of genetic biomarkers have been discovered through large-scale genome wide association studies (GWASs) in Alzheimer’s disease (AD), such as *APOE, TOMM40* and *CLU*. Despite this significant progress, existing genetic findings are largely passengers not directly involved in the driver events, which presents challenges for replication and translation into targetable mechanisms. In this paper, leveraging the protein interaction network, we proposed a modularity-constrained Lasso model to jointly analyze the genotype, gene expression and protein expression data. With a prior network capturing the functional relationship between SNPs, genes and proteins, the newly introduced penalty term maximizes the global modularity of the subnetwork involving selected markers and encourages the selection of multi-omic markers with dense functional connectivity, instead of individual markers. We applied this new model to the real data in ROS/MAP cohort for discovery of biomarkers related to cognitive performance. A functionally connected subnetwork involving 276 multi-omic biomarkers, including SNPs, genes and proteins, were identified to bear predictive power. Within this subnetwork, multiple trans-omic paths from SNPs to genes and then proteins were observed, suggesting that cognitive performance can be potentially affected by the genetic mutations due to their cascade effect on the expression of downstream genes and proteins.

## Introduction

Alzheimer’s disease (AD) is the most common form of brain dementia characterized by the gradual loss of memory and other cognitive function. With rapidly increasing aging population, AD is drawing more and more attention in the United States and around the world [1]. Unfortunately, the underlying mechanism of AD remains largely unknown and no clinically validated drug is available for disease treatment and prevention. Although recent large-scale genome wide association studies (GWASs) have led to discovery of many genetic markers associated with AD, such as *APOE, TOMM40* and *CLU*, replicability of existing findings and their translation into targetable mechanisms related to disease pathogenesis remain a challenge. Identification of novel biomarkers or functionally validating existing biomarkers becomes increasingly important for discovery of new potential future therapeutic targets.

Recently, there is a substantial increase in AD multi-omic data. Example projects include the Alzheimer’s Disease Neuroimaging Initiative (ADNI) [2] and the Religious Orders Study and Memory and Aging Project (ROSMAP) [3]. Instead of limiting their perspective to a single -omics layer, these data collections create a molecular landscape spanning the genome, transcriptome, proteome and metabolome. Coupling with systems biology networks (e.g., protein-protein interaction (PPI) network), these data provides a valuable resource with rich content and opens numerous opportunities for more comprehensive analyses of AD. These multi-omics data has been increasingly recognized to be a potential key enabler of novel biomarker discovery [4, 5]. It not only allows us to examine the disease from different-omics layers, but also provides insights into their interactions which is critical for translation of genetic findings into targetable mechanism.

Despite this great potential, the power of multi-omic data has not been fully unleashed. Much research effort of existing studies has been on single type of -omics data without acknowledging the interconnections between –omics layers. This shortcoming is largely due to the limited availability of computational methods that are sufficiently powerful and comprehensive enough to handle the high dimensionality and heterogeneity of multimic data. In addition, major findings generated from current –omics studies have been largely restricted to relatively simple patterns. They are mostly individual biomarkers, possibly without functional interactions, which presents difficulties to validate these findings and to relate them to downstream biology [6, 7]. To address this problem, some recent studies propose to seek common genetic markers with evidence from more than one -omics layer [8–10], which are expected to more reliable for further experimental validation. However, this simple overlap strategy may be too stringent as -omic features in different layers are not completely mapped in a one-to-one relationship.

In this paper, leveraging the functional interaction network in REACTOME, we propose a modularity-constrained Lasso model to jointly analyze genotype, gene expression and protein expression data. We aim to identify a set of SNPs, genes and proteins as biomarkers, forming a subnetwork with functional connections cutting across different-omics layers. Compared to individual mutations, genes or proteins identified using traditional methods, such connected pattern can help improve not only the reliability of identified biomarkers, but also their replicability and interpretability.

## Method

### Study cohort

All the data analyzed in the present report were obtained from the Religious Orders Study (ROS) and Memory and Aging Project (MAP). It was launched by Rush University to build a cohort from religious communities to measure the progression of amnestic mild cognitive impairment (MCI, a prodromal stage of AD) and early probable AD. The combined ROS/MAP cohort includes around 600 participants under age 90, which constitute a very rich repository of multi-modal data including GWAS data, whole genome sequencing (WGS) data, cognitive, behavioral and clinical data. The more detailed description could be found in [3]. In this paper, GWAS genotype data and quality controlled RNA-Seq gene expression and protein expression data collected from prefrontal cortex tissue in the brain were downloaded. To perform the proposed joint analysis, only subjects with all three types of -omics data were included. In total, we have 262 subjects (115 healthy controls (HC), 67 mild cognitive impairment (MCI), 80 AD patients) with full set of genotype, RNA-seq gene expression and proteomic data. The detailed demographic information can be found in Table 1.

**Table 1.** Demographic information of the ROS/MAP participants included in this study.

|  |  |  |  |
| --- | --- | --- | --- |
| Dignosis | HC | MCI | AD |
| Subject Number | 115 | 67 | 80 |
| ROS/MAP | 69/46 | 27/40 | 40/40 |
| Male/Female | 51/64 | 28/39 | 31/49 |
| Education(mean± std.) | 16.9 ± 3.5 | 16.6 ± 3.3 | 16.9 ± 3.8 |
| Age(mean± std.) | 83.0 ± 4.7 | 85.0 ± 4.2 | 86.3 ± 3.7 |

### GWAS genotype data preparation

ROS/MAP samples were genotypes on the Affymetrix GeneChip 6.0 platform [11]. We performed sample and SNP quality control procedures on GWAS data (SNP call rate<95%, Hardy-Weinberg equilibrium test p<10^−6^ in controls, and frequency filtering (MAF<1%) were performed. After performing the standard quality control procedures for genetic markers and subjects, only non-Hispanic Caucasian participants were selected by clustering with CEU (Utah residents with Northern and Western European ancestry from the CEPH collection) + TSI (Toscani in Italia) populations using HapMap 3 genotype data and the multidimensional scaling (MDS) analysis [12]. Un-genotyped SNPs were imputed using MaCH and the 1000 Genomes Project as a reference panel [13].

### RNA-Seq gene expression preparation

RNA-Seq gene expression data in the ROS/MAP cohort were collected from the prefrontal cortex tissue in the brain. The RNA-Seq data were recently reprocessed in parallel with other AMP-AD RNAseq datasets, and this second version of the data were downloaded for our subsequent analysis. The input data for the RNAseq reprocessing effort was aligned reads in bam files that were converted to fastq using the Picard SamToFastq function. Fastq files were re-aligned to the reference genome using STAR with twopassMode set as Basic. Gene counts were computed for each sample by STAR by setting quantMode as GeneCounts. These gene level counts further went through normalized and adjusted to remove the effects of relevant factors such as age, gender, education, batch, RNA integrity number (RIN) and post moterm interval (PMI). Detailed reprocessing and normalization steps can be found in the AMP-AD knowledge portal (https://www.synapse.org/#!Synapse:syn9702085/).

### Protein expression data preparation

SRM proteomics was performed using frozen tissue from dorsolateral prefrontal cortex (DLPFC). The samples were prepared for LC-SRM analysis using standard protocol as described in [14, 15]. All the data were manually inspected to ensure correct peak assignment and peak boundaries.The abundance of endogenous peptides was quantified as a ratio to spiked-in synthetic peptides containing stable heavy isotopes. The “light/heavy” ratios were log2 transformed and shifted such that median log2-ratio is zero. Normalization adjusted for differences in protein amounts between the samples. During that normalization, we shifted the log2-ratios for each sample to make sure the median is set at zero. Detailed processing steps can be found in the AMP-AD knowledge portal (https://www.synapse.org/#!Synapse:syn8456629). Using the regression weights derived from the healthy control participants, peptide abundance data were further adjusted to remove the effects of the age at death, gender, education, PMI and batch.

### Selection of SNPs, genes and proteins

We focused our analysis on a set of SNPs, genes and proteins with known functional connections. Though we have genome-wide genotype and transcriptome-wide gene expression data available in the ROS/MAP cohort, only a limited number of proteins are measured and form a bottleneck for the joint -omics data analysis. To address this problem, we used these proteins as seeds to select a subset of SNPs and genes for subsequent analysis. As shown in Fig 1, in the proteomic level, abundance level of 186 peptides, from 126 unique genes, were measured in the ROS/MAP project. When mapped to the functional interaction network in REACTOME [16], where all protein interactions were manually curated from pathways with directionality information, these genes are found to interact with 954 genes. After excluding those without gene ID, totally 743 genes with RNA-seq data were included in the transcriptomic level. In the genomic level, SNPs located on the upstream of these genes (boundary: 5K) were extracted. To ensure the functional connection of selected SNPs and their downstream genes, we included only SNPs significantly affecting the transcrition factor binding activity, as shown in SNP2TFBS database [17]. This relationship between SNPs, genes and proteins/peptides are used as the trans-omic network to guide the search of functionally connected biomarkers in the subsequent analysis.

**Fig 1.**
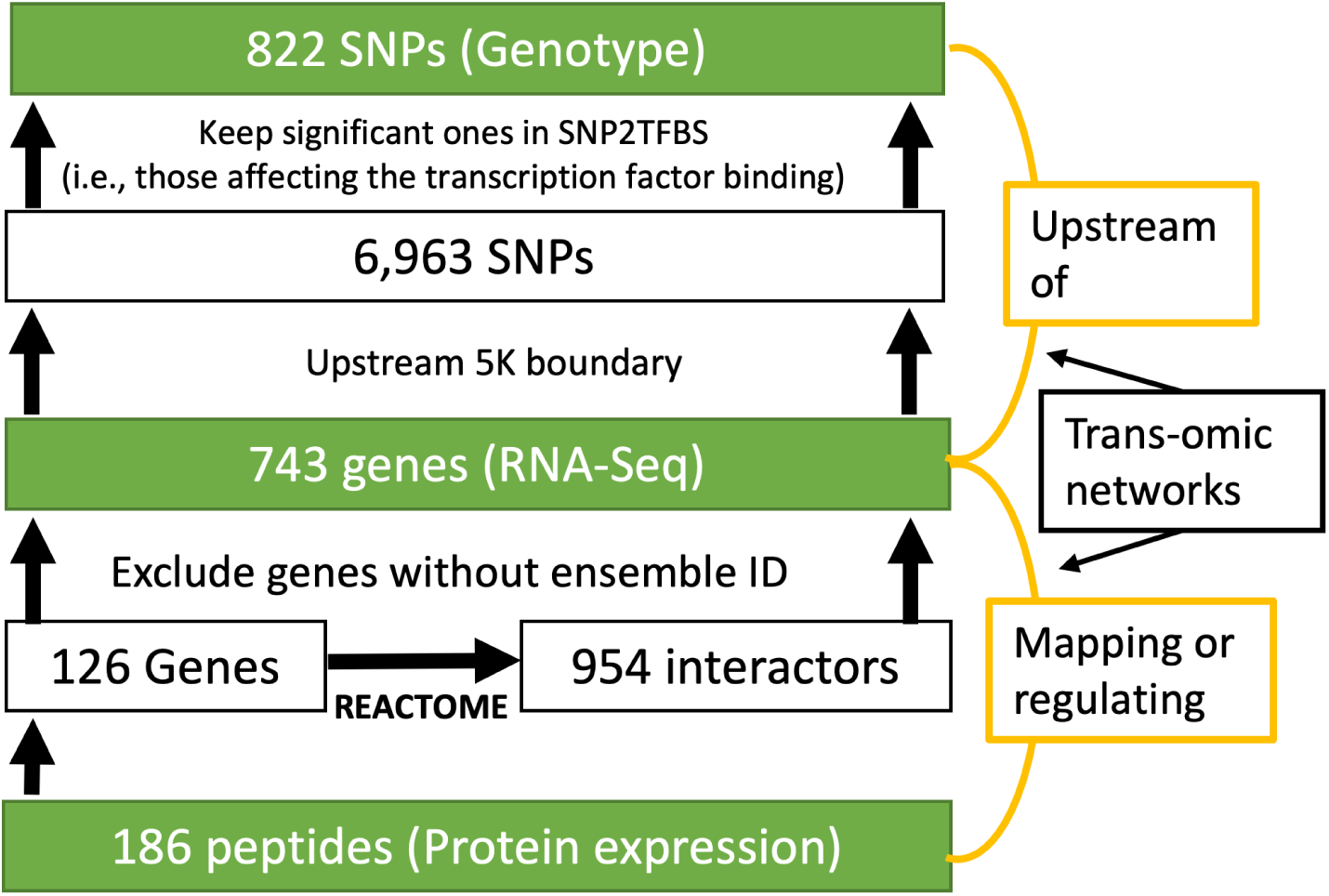
The selection of SNPs in upstream 5K boundary for each gene.

### Memory outcomes

In the ROS/MAP project, cognitive performance of participants was estimated through the mini mental state examination, a standardized screening measure for collecting 30 items in related with dementia [18, 19]. This score ranges from 0 to 30, and is scaled to quantify the severity of dementia. In this study, we use this memory test score as the AD quantitative trait for discovery of functionally connected biomarkers. Using the regression weights derived from the healthy control participants, the memory score is adjusted to remove the effect of sex, education and age.

### Modularity-constrained Lasso

Throughout this section, we write matrices as boldface uppercase letters and vectors as boldface lowercase letters. Given a matrix **M** = (*m*_*ij*_), its *i*-th row and *j*-th column are denoted as **m**^*i*^ and **m**_*j*_ respectively. Let **X** = [***x***_1_, ***x***_2_, …, ***x***_*n*_]^*T*^ be the multi-omic features as predictors and **y** = [*y*_1_, *y*_2_, …, *y*_*n*_]^*T*^ be the disease quantitative trait as outcome (i.e., cognitive performance). Here, ***x***_*j*_ ⊆ ℝ^*p*^ is a concatenated vector of genotype, gene expression and protein expression data for *j*-th subject.

The least absolute shrinkage and selection operator (Lasso) is a shrinkage and selection method for linear regression [20]. It minimizes the usual sum of squared errors with a bound on the sum of the absolute values of the coefficients, which is also known as L1 norm (Eq. 1).

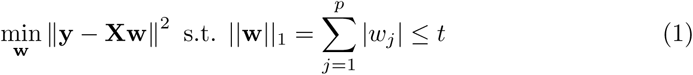

With this constraint, Lasso aims to minimize the number of selected features, which significantly improved the interpretability of results compared to traditional linear regression, where almost all features are considered to be outcome-relevant with non-zero weight. However, when dealing with a group of highly correlated features, L1 norm penalty will result in a random selection. In this case, multiple runs of Lasso on the same set of data will possibly generate different set of selected features, which presents challenges for replicating and interpreting the results. To address this problem, several groups proposed to explicitly incorporate the correlation structure into the sparse prediction model and encourage the selection/exclusion of all highly correlated features [21–24]. Among those is GraphNet, where a graph **G** ⊆ ℝ^*p*×*p*^ indicating the correlation structure between predictors is used as a priori to guide the feature selection (Eq. 2) [24]. Here, **L** is the corresponding Laplacian matrix of graph **G**. However, GraphNet only takes account into local topology information with a focus on pairwise similarity. For multi-omic biomarker discovery, using this penalty can not guarantee the selected features are densely connected in the prior network.

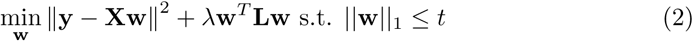

In this paper, we propose a new modularity-constrained Lasso which leverages the global network property to encourage the selection of a sub-network module rather than individual markers scattered in the prior network. Given the trans-omic network capturing the functional interaction between SNPs, genes and proteins, we formulate it as a graph and its corresponding adjacency matrix is denoted as **G** ⊆ ℝ^*p*×*p*^. **B** is the modularity matrix [25], where 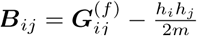. It evaluates whether the number of links is significantly more than expected. *h*_*i*_ and *h*_*j*_ are the degrees of the *i*-th and h node in the network, and *m* is the total number of links in the network. To impose a modular structure in the identified biomarkers, we propose a new penalty term as *P*_*M*_ (**w, B**) =< **w**^*T*^ **w, B** *>*, inspired by the module identification problem [26, 27]. Here, *<>* is the Frobenius inner product defined by *< A, B >*= *tr*(*A*^*T*^ *B*). Maximizing the Frobenius inner product between **w**^*T*^ **w** and the modularity matrix **B** encourages the selection of features with dense functional connections in the prior multi-omic network.

Taken together, our new modularity-constrained Lasso objective is formulated as in Eq. 3.

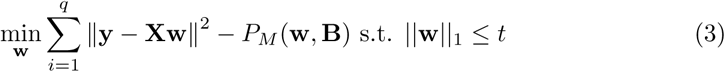

Here, *λ* and *t* are the parameters that control and balance the contribution from two regularization terms. Note that the objective function in Eq. 3 is not convex because the modularity matrix **B** used in *P*_*M*_ (**w, B**) =< **w**^*T*^ **w, B** *>* is indefinite. To make **B** negative-definite, we introduced an auxiliary function where **B** is replaced by **B** − *λ*_*B*_**I** and *λ*_*B*_ is the absolute maximum eigenvalue of **B**. Eq. 3 can be easily solved by obtaining a closed form solution without L1 constraint, followed by soft-threshold method [20].

## Results

### Performance comparison between M-Lasso and G-Lasso

In this section, we denote our modularity-constrained Lasso as M-Lasso and GraphNet-constrained Lasso as G-Lasso. For both methods, nested 5-fold cross validation (CV) procedure was applied to tune the parameters based on root mean squared error (RMSE) and the portion of different diagnosis groups was kept the same in different folds. As can be observed, the major difference between Eq. 2 and Eq. 3 is the penalty term and they have the same set of parameters. The parameters are tuned with the range set from [10^−6^ 10^−5^ 10^−4^ 10^−3^ 10^−2^ 10^−1^ 10^0^ 10^1^ 10^2^]. For fair comparison, both methods were evaluated using the same partition of subjects during the cross validation procedure.

Shown in Table 2 is the root mean square error estimated by M-Lasso and G-Lasso on test data set across five folds. As we can observe, M-Lasso consistently outperforms G-Lasso with smaller prediction error over all 5 folds.

**Table 2.** Performance comparison on test set between M-Lasso and G-Lasso (RMSE).

|  | Fold 1 | Fold 2 | Fold 3 | Fold 4 | Fold 5 | Mean |
| --- | --- | --- | --- | --- | --- | --- |
| M-Lasso | 0.853 | 1.082 | 0.905 | 0.959 | 0.874 | 0.935 |
| G-Lasso | 0.876 | 1.127 | 1.015 | 1.085 | 1.415 | 1.104 |

For feature selection, M-Lasso identified around 600 -omics features, including SNPs, genes and proteins, to be predictive of cognitive performance, while G-Lasso only identified a handful of them (i.e., less than 20 for all 5 folds). When mapped to the prior functional connectivity network, markers identified by G-Lasso scatters across the network with few connections, which suggests that the local topology information used in GraphNet penalty is not strong enough to form subnetwork structure among identified biomarkers. For M-Lasso, -omics biomarkers identified are largely connected to each other in the prior network. Take the result from one fold as example, 650 -omics features were selected, including 255 SNPs, 339 genes and 56 proteins. The largest connected network component involves 276 -omics features with 366 edges (Fig. 2). The rest of the multi-omic markers identified in M-Lasso mostly form small connected components, ranging in size from 2 to 50. These features are found predictive yet not well functionally connected, possibly due to the fact that they are false positives or their functional connections have not been previously studied yet. In the subsequent part, we focus on the multi-omic biomarkers in the largest connected component, which are both predictive of cognitive performance and functionally connected with evidence from prior knowledge.

**Fig 2.**
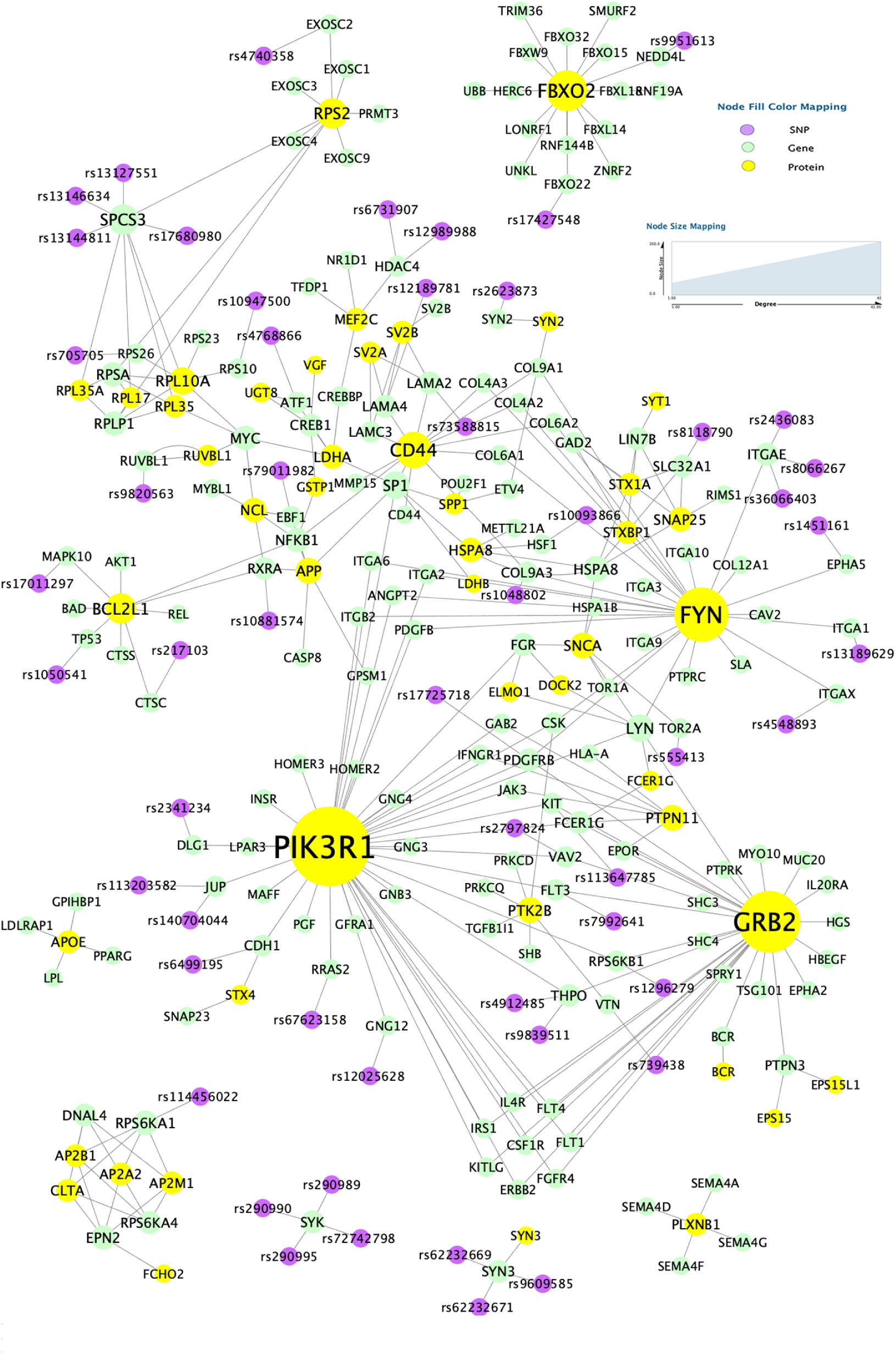
Top 7 connected components with biomarkers identified using M-Lasso mapped to prior network.

### Functionally connected multi-omic biomarkers

Shown in Fig. 2 are the top 7 connected components obtained after mapping 650 -omics features back to the prior network. Size of each node is made proportional to their degree. It can be easily observed that there are multiple trans-omic paths from SNPs to genes and then proteins. Note that these SNPs are located upstream of their connected genes and has significant effect on the transcription factor binding activity. Thus, these SNPs are very likely to have an influence on the expression of their connected genes. Also, the functional interaction between genes and proteins are curated from REACTOME pathways with direction information. Therefore, genes have a regulatory role toward the expression of their connected proteins in the prior network. Taken together, these trans-omic paths suggest that cognitive performance can be potentially affected by the genetic mutations (i.e., SNPs) due to their cascade effect on the expression of downstream genes, which further regulate the protein expression. We further examined all 28 SNPs involved in the largest connected component in BRAINEAC database. 25 of them were found to be expression quantitative locus (eQTLs) in the prefrontal cortex tissue, which gives further support to our discovery of trans-omic paths as biomarkers.

For the largest connected components, we further performed network analysis using NetworkAnalyzer in Cytoscape [28] and identified the -omic biomarkers with top centrality values, such as degree, betweenness and closeness (Table 3). Top nodes by degree in this subnetwork included proteins *PIK3R1, FYN, CD44* and *RPS2*, and genes *GRB2, FBXO2, EP300, SV2A* and *SPCS3*. Most hub nodes are also found to have the top centrality value in betweenness, closeness and clustering coefficient, such as *PIK3R1, FYN*, and *EP300*. Majority of these genes and proteins have been previously reported in association with AD. For example, *PIK3R1* encodes the regulatory subunit of the phosphoinositide-3-kinase protein complex PI3Ks, which are known to play a key role in insulin signaling. Results from recent studies start to show evidence of intrinsic insulin resistance inside AD brains [29]. The hub gene *EP300* encodes the enzyme histone acetyltransferase *p300* or E1A-associated protein *p300*, also known as *EP300* or *P300*. This enzyme functions as histone acetyltransferase that regulates transcription of genes via chromatin remodeling. Findings from multiple studies have suggested the potential of *P300* to act as a biomarker for dementia assessment and monitoring AD. Meta-analysis of *P300* amplitude and latency reveals useful information about the early stages of AD [30, 31]. In addition, both *GRB2* and *FBXO2* were found to interact with *APP*, a well-known gene related to AD. *GRB2* interacts with *APP* requiring phosphorylation of APP at Tyr-682 [32]. This could lead to the activation of the MAPK pathway, since *GRB2* are known to link growth factor receptors to signaling pathways, such as MAPK and PI3K, and participate in oncogenic proliferation, neuronal development, cell differentiation, and apoptosis [33–38] The hub gene *FBXO2* participates in *APP* processing by promoting degradation of *APP* cleaving *β*-secretase [39].

**Table 3.** For 7 network features, nodes with the top highest value in modularity enrichment analysis.

|  |  |
| --- | --- |
| Degree | *PIK3R1*, ***GRB2***, *FYN*, ***FBXO2***, *CD44*, *HSPA8*, ***EP300***, *RPS2*, ***SV2A***, ***SPCS3*** |
| Average Shortest Path Length | *PIK3R1*, ***SP1***, *FYN*, ***EP300***, ***LYN***, *LDHA*, ***PDGFRB***, *APP*, ***GRB2***, ***FCER1G*** |
| Betweenness | *PIK3R1*, ***SP1***, ***EP300***, *FYN*, ***GRB2***, ***MYC***, *LDHA*, *CD44*, ***RPL10A***, *HSPA8* |
| Closeness | *PIK3R1*, ***SP1***, *FYN*, ***EP300***, ***LYN***, *LDHA*, ***PDGFRB***, *APP*, ***GRB2***, ***FCER1G*** |
| Clustering Coefficient | ***FCER1G***, ***LYN***, *GRB2*, *PIK3R1* |

rs- IDs: SNPs; Bold: genes; The rest: proteins.

### Pathway enrichment analysis

For 168 genes and 37 proteins in the largest connected subnetwork, we performed pathway enrichment analysis using ClueGO based on the Kyoto Encyclopedia of Genes and Genomes (KEGG) database [40, 41]. In total, 74 pathways were found to be significantly enriched by our gene/protein set, with Bonferroni corrected term p-value smaller than 5% (P ≤0.05). Shown in Table 4 was the top 20 enriched KEGG pathways with smallest p values after correction. The top hit is *PI3K* -*Akt* signaling pathway, a major mediator of effects of insulin. Two recent studies have found a significant correlation between peripheral insulin resistance and brain A*β* levels as measured by Pittsburgh compound B-positron emission tomography (PiB-PET) [42, 43]. The impaired insulin-PI3K-Akt signaling observed in the AD brain has led to clinical trials studying whether the enhancement of this pathway using intranasal insulin (IN) treatment is beneficial [44]. Other enriched pathways that are previously reported with a key role in AD include Focal adhesion [45], Ras signaling pathway [46], ECM-receptor interaction [47], MAPK signaling pathway [48], Rap1 signaling pathway [49], etc. In addition, we observe many of the top enriched pathways are related to cancer, such as *PI3K* -*Akt* signaling pathway, Prostate cancer and small lung cancer. This finding provide support to the hypothesis of shared pathological mechanism between cancer and AD [50–53].

**Table 4.** Top enriched KEGG pathways by the gene and protein markers in the largest subnetwork.

| Pathway | Number of markers in the pathway | Total number of genes in the pathway | p-value | Corrected p-value |
| --- | --- | --- | --- | --- |
| *PI3K-Akt* signaling pathway | 84 | 354 | 5.78E-42 | 9.99E-40 |
| Focal adhesion | 59 | 199 | 5.89E-35 | 1.01E-32 |
| Pathways in cancer | 86 | 530 | 1.25E-29 | 2.14E-27 |
| Ras signaling pathway | 52 | 232 | 2.20E-24 | 3.73E-22 |
| Human papillomavirus infection | 55 | 330 | 2.88E-19 | 4.87E-17 |
| *ECM*-receptor interaction | 28 | 82 | 9.34E-19 | 1.57E-16 |
| *MAPK* signaling pathway | 51 | 295 | 1.34E-18 | 2.24E-16 |
| Kaposi sarcoma-associated herpesvirus infection | 39 | 186 | 2.40E-17 | 3.98E-15 |
| Rap1 signaling pathway | 41 | 206 | 2.48E-17 | 4.09E-15 |
| Human cytomegalovirus infection | 42 | 225 | 1.14E-16 | 1.87E-14 |
| Prostate cancer | 27 | 97 | 1.41E-15 | 2.30E-13 |
| Chronic myeloid leukemia | 24 | 76 | 2.38E-15 | 3.85E-13 |
| *ErbB* signaling pathway | 25 | 85 | 3.99E-15 | 6.42E-13 |
| Small cell lung cancer | 25 | 93 | 4.04E-14 | 6.46E-12 |
| Phospholipase D signaling pathway | 31 | 148 | 5.71E-14 | 9.08E-12 |
| Chemokine signaling pathway | 35 | 190 | 7.46E-14 | 1.18E-11 |
| Proteoglycans in cancer | 36 | 201 | 7.87E-14 | 1.24E-11 |
| Neurotrophin signaling pathway | 27 | 119 | 3.39E-13 | 5.28E-11 |
| Relaxin signaling pathway | 28 | 130 | 4.84E-13 | 7.50E-11 |
| JAK-STAT signaling pathway | 30 | 162 | 4.38E-12 | 6.74E-10 |

## Conclusion

In this study, we proposed a new modularity-constrained Lasso model to jointly analyze the genotype, RNA-Seq gene expression and protein expression data. The newly introduced penalty term maximizes the global modularity of selected biomarkers in the prior network and encourages the selection of multi-omic biomarkers forming network modules. Compared to the GraphNet penalty that enforces local pairwise similarity, modularity-based penalty helps identify more biomarkers with significantly improved functional connectivity. In particular, we found that some biomarkers form trans-omic paths from SNP to gene and then protein, suggesting the potential cascade effect of genotype on the downstream transcriptome and proteome level. To the best of our knowledge, this is the first study that explored the potential of functional multi-omic subnetworks as biomarkers in AD.

Despite the promising findings, this study has multiple limitations. First, only one disease quantitative trait is used as outcome in the prediction model. Considering the potential bias introduced from data collection procedure, the biomarkers and their functional connectivity network identified here may not reflect the optimal pattern. Incorporating multiple correlated outcomes and performing a multitask prediction will possibly help improve the performance. Second, our proposed model is not capable of handling the missing data problem. Each subject has to have all the –omics data to be included in the analysis. Therefore, many subjects with missing data in one or more –omics layers are inevitably excluded and only a small portion of the big –omics data is utilized. Future efforts are warranted to further improve this model.

## Acknowledgments

This research was supported by NIH grants R21 AG066135, R01 EB022574, R01 AG019771, P30 AG010133, NSF CRII 1755836 and Indiana University Collaborative Research Grant (IUCRG). This project was also funded, in part, with support from the Indiana Clinical and Translational Sciences Institute funded, in part by Grant Number UL1TR001108 from the National Institutes of Health, National Center for Advancing Translational Sciences, Clinical and Translational Sciences Award.

## References

1. Organization WH, et al. Global Health Estimates 2016: Deaths by Cause. Age, Sex, by Country and by Region. 2000;2016:2018.

2. Mueller SG, Weiner MW, Thal LJ, Petersen RC, Jack C, Jagust W, et al. The Alzheimer’s disease neuroimaging initiative. Neuroimaging Clinics. 2005;15(4):869–877.

3. A Bennett D, A Schneider J, Arvanitakis Z, S Wilson R. Overview and findings from the religious orders study. Current Alzheimer Research. 2012;9(6):628–645.

4. Hasin Y, Seldin M, Lusis A. Multi-omics approaches to disease. Genome biology. 2017;18(1):83.

5. Huang S, Chaudhary K, Garmire LX. More is better: recent progress in multiomics data integration methods. Frontiers in genetics. 2017;8:84.

6. Visscher PM, Brown MA, McCarthy MI, Yang J. Five years of GWAS discovery. The American Journal of Human Genetics. 2012;90(1):7–24.

7. Edwards SL, Beesley J, French JD, Dunning AM. Beyond GWASs: illuminating the dark road from association to function. The American Journal of Human Genetics. 2013;93(5):779–797.

8. Whitaker JW, Boyle DL, Bartok B, Ball ST, Gay S, Wang W, et al. Integrative omics analysis of rheumatoid arthritis identifies non-obvious therapeutic targets. PLoS One. 2015;10(4):e0124254.

9. Zhang JG, Tan LJ, Xu C, He H, Tian Q, Zhou Y, et al. Integrative analysis of transcriptomic and epigenomic data to reveal regulation patterns for BMD variation. PloS one. 2015;10(9):e0138524.

10. Lin D, Zhang J, Li J, He H, Deng HW, Wang YP. Integrative analysis of multiple diverse omics datasets by sparse group multitask regression. Frontiers in cell and developmental biology. 2014;2:62.

11. De Jager PL, Shulman JM, Chibnik LB, Keenan BT, Raj T, Wilson RS, et al. A genome-wide scan for common variants affecting the rate of age-related cognitive decline. Neurobiology of aging. 2012;33(5):1017–e1.

12. Horgusluoglu-Moloch E, Nho K, Risacher SL, Kim S, Foroud T, Shaw LM, et al. Targeted neurogenesis pathway-based gene analysis identifies ADORA2A associated with hippocampal volume in mild cognitive impairment and Alzheimer’s disease. Neurobiology of aging. 2017;60:92–103.

13. Nho K, Corneveaux J, Kim S, Lin H, Risacher S, Shen L, et al. Whole-exome sequencing and imaging genetics identify functional variants for rate of change in hippocampal volume in mild cognitive impairment. Molecular psychiatry. 2013;18(7):781.

14. Petyuk VA, Qian WJ, Smith RD, Smith DJ. Mapping protein abundance patterns in the brain using voxelation combined with liquid chromatography and mass spectrometry. Methods. 2010;50(2):77–84.

15. Andreev VP, Petyuk VA, Brewer HM, Karpievitch YV, Xie F, Clarke J, et al. Label-free quantitative LC–MS proteomics of Alzheimer’s disease and normally aged human brains. Journal of proteome research. 2012;11(6):3053–3067.

16. Fabregat A, Jupe S, Matthews L, Sidiropoulos K, Gillespie M, Garapati P, et al. The reactome pathway knowledgebase. Nucleic acids research. 2017;46(D1):D649–D655.

17. Kumar S, Ambrosini G, Bucher P. SNP2TFBS–a database of regulatory SNPs affecting predicted transcription factor binding site affinity. Nucleic acids research. 2016;45(D1):D139–D144.

18. Folstein MF, Folstein SE, McHugh PR. “Mini-mental state”: a practical method for grading the cognitive state of patients for the clinician. Journal of psychiatric research. 1975;12(3):189–198.

19. Folstein MF, Robins LN, Helzer JE. The mini-mental state examination. Archives of general psychiatry. 1983;40(7):812–812.

20. Tibshirani R. Regression shrinkage and selection via the lasso. Journal of the Royal Statistical Society: Series B (Methodological). 1996;58(1):267–288.

21. Jacob L, Obozinski G, Vert JP. Group lasso with overlap and graph lasso. In: Proceedings of the 26th annual international conference on machine learning. ACM; 2009. p. 433–440.

22. Yuan L, Liu J, Ye J. Efficient methods for overlapping group lasso. In: Advances in Neural Information Processing Systems; 2011. p. 352–360.

23. Kim S, Xing EP, et al. Tree-guided group lasso for multi-response regression with structured sparsity, with an application to eQTL mapping. The Annals of Applied Statistics. 2012;6(3):1095–1117.

24. Grosenick L, Klingenberg B, Katovich K, Knutson B, Taylor JE. Interpretable whole-brain prediction analysis with GraphNet. NeuroImage. 2013;72:304–321.

25. Newman ME. Modularity and community structure in networks. Proceedings of the national academy of sciences. 2006;103(23):8577–8582.

26. Hildebrand R. Identification of community structure in networks with convex optimization. arXiv preprint 08061896. 2008;.

27. Chan YK, Yeung DY. A convex formulation of modularity maximization for community detection. In: Twenty-Second International Joint Conference on Artificial Intelligence; 2011.

28. Shannon P, Markiel A, Ozier O, Baliga NS, Wang JT, Ramage D, et al. Cytoscape: a software environment for integrated models of biomolecular interaction networks. Genome research. 2003;13(11):2498–2504.

29. Arnold SE, Arvanitakis Z, Macauley-Rambach SL, Koenig AM, Wang HY, Ahima RS, et al. Brain insulin resistance in type 2 diabetes and Alzheimer disease: concepts and conundrums. Nature Reviews Neurology. 2018;14(3):168.

30. Polich J, Ladish C, Bloom FE. P300 assessment of early Alzheimer’s disease. Electroencephalography and Clinical Neurophysiology/Evoked Potentials Section. 1990;77(3):179–189.

31. Hedges D, Janis R, Mickelson S, Keith C, Bennett D, Brown BL. P300 amplitude in Alzheimer’s disease: a meta-analysis and meta-regression. Clinical EEG and neuroscience. 2016;47(1):48–55.

32. Roncarati R, Šestan N, Scheinfeld MH, Berechid BE, Lopez PA, Meucci O, et al. The *γ*-secretase-generated intracellular domain of *β*-amyloid precursor protein binds Numb and inhibits Notch signaling. Proceedings of the National Academy of Sciences. 2002;99(10):7102–7107.

33. Russo C, Dolcini V, Salis S, Venezia V, Zambrano N, Russo T, et al. Signal transduction through tyrosine-phosphorylated C-terminal fragments of amyloid precursor protein via an enhanced interaction with Shc/Grb2 adaptor proteins in reactive astrocytes of Alzheimer’s disease brain. Journal of Biological Chemistry. 2002;277(38):35282–35288.

34. Cattaneo E, Pelicci PG. Emerging roles for SH2/PTB-containing Shc adaptor proteins in the developing mammalian brain. Trends in neurosciences. 1998;21(11):476–481.

35. Yokote K, Mori S, Hansen K, McGlade J, Pawson T, Heldin CH, et al. Direct interaction between Shc and the platelet-derived growth factor beta-receptor. Journal of Biological Chemistry. 1994;269(21):15337–15343.

36. Saucier C, Papavasiliou V, Palazzo A, Naujokas MA, Kremer R, Park M. Use of signal specific receptor tyrosine kinase oncoproteins reveals that pathways downstream from Grb2 or Shc are sufficient for cell transformation and metastasis. Oncogene. 2002;21(12):1800.

37. Napoli C, Martin-Padura I, De Nigris F, Giorgio M, Mansueto G, Somma P, et al. Deletion of the p66Shc longevity gene reduces systemic and tissue oxidative stress, vascular cell apoptosis, and early atherogenesis in mice fed a high-fat diet. Proceedings of the National Academy of Sciences. 2003;100(4):2112–2116.

38. Dankort D, Maslikowski B, Warner N, Kanno N, Kim H, Wang Z, et al. Grb2 and Shc adapter proteins play distinct roles in Neu (ErbB-2)-induced mammary tumorigenesis: implications for human breast cancer. Molecular and cellular biology. 2001;21(5):1540–1551.

39. Atkin G, Hunt J, Minakawa E, Sharkey L, Tipper N, Tennant W, et al. F-box only protein 2 (Fbxo2) regulates amyloid precursor protein levels and processing. Journal of Biological Chemistry. 2014;289(10):7038–7048.

40. Bindea G, Mlecnik B, Hackl H, Charoentong P, Tosolini M, Kirilovsky A, et al. ClueGO: a Cytoscape plug-in to decipher functionally grouped gene ontology and pathway annotation networks. Bioinformatics. 2009;25(8):1091–1093.

41. Kanehisa M, Goto S. KEGG: kyoto encyclopedia of genes and genomes. Nucleic acids research. 2000;28(1):27–30.

42. Willette AA, Johnson SC, Birdsill AC, Sager MA, Christian B, Baker LD, et al. Insulin resistance predicts brain amyloid deposition in late middle-aged adults. Alzheimer’s & dementia. 2015;11(5):504–510.

43. Ekblad LL, Johansson J, Helin S, Viitanen M, Laine H, Puukka P, et al. Midlife insulin resistance, APOE genotype, and late-life brain amyloid accumulation. Neurology. 2018;90(13):e1150–e1157.

44. Gabbouj S, Ryhänen S, Marttinen M, Wittrahm R, Takalo M, Kemppainen S, et al. Altered insulin signaling in Alzheimer’s disease brain–special emphasis on PI3K-Akt pathway. Frontiers in neuroscience. 2019;13:629.

45. Caltagarone J, Jing Z, Bowser R. Focal adhesions regulate A*β* signaling and cell death in Alzheimer’s disease. Biochimica et Biophysica Acta (BBA)-Molecular Basis of Disease. 2007;1772(4):438–445.

46. Kirouac L, Rajic AJ, Cribbs DH, Padmanabhan J. Activation of Ras-ERK signaling and GSK-3 by amyloid precursor protein and amyloid beta facilitates neurodegeneration in Alzheimer’s disease. Eneuro. 2017;4(2).

47. Kerrisk ME, Cingolani LA, Koleske AJ. ECM receptors in neuronal structure, synaptic plasticity, and behavior. In: Progress in brain research. vol. 214. Elsevier; 2014. p. 101–131.

48. Zhu X, Lee Hg, Raina AK, Perry G, Smith MA. The role of mitogen-activated protein kinase pathways in Alzheimer’s disease. Neurosignals. 2002;11(5):270–281.

49. Dumbacher M, Van Dooren T, Princen K, De Witte K, Farinelli M, Lievens S, et al. Modifying Rap1-signalling by targeting Pde6δ is neuroprotective in models of Alzheimer’s disease. Molecular neurodegeneration. 2018;13(1):50.

50. W Nixon D. The inverse relationship between cancer and Alzheimer’s Disease: A possible mechanism. Current Alzheimer Research. 2017;14(8):883–893.

51. Musicco M, Adorni F, Di Santo S, Prinelli F, Pettenati C, Caltagirone C, et al. Inverse occurrence of cancer and Alzheimer disease: a population-based incidence study. Neurology. 2013;81(4):322–328.

52. Roe C, Behrens M, Xiong C, Miller J, Morris J. Alzheimer disease and cancer. Neurology. 2005;64(5):895–898.

53. Majd S, Power J, Majd Z. Alzheimer’s Disease and Cancer: When Two Monsters Cannot Be Together. Frontiers in neuroscience. 2019;13.

